# Non-prime- and Prime-side Profiling of Pro-Pro Endopeptidase Specificity Using Synthetic Combinatorial Peptide Libraries and Mass Spectrometry

**DOI:** 10.1101/2024.03.15.585006

**Authors:** Bart Claushuis, Robert A. Cordfunke, Arnoud H. de Ru, Jordy van Angeren, Ulrich Baumann, Peter A. van Veelen, Manfred Wuhrer, Jeroen Corver, Jan W. Drijfhout, Paul J. Hensbergen

## Abstract

A group of bacterial proteases, the Pro-Pro endopeptidases (PPEPs), possess the unique ability to hydrolyze proline-proline bonds in proteins. Since a protease’s function is largely determined by its substrate specificity, methods that can extensively characterize substrate specificity are valuable tools for protease research. Previously, we achieved an in-depth characterization of PPEP prime-side specificity. However, PPEP specificity is also determined by the non-prime-side residues in the substrate.

To gain a more complete insight into the determinants of PPEP specificity, we characterized the non-prime- and prime-side specificity of various PPEPs using a combination of synthetic combinatorial peptide libraries and mass spectrometry. With this approach, we deepened our understanding of the P3-P3’ specificities of PPEP-1 and PPEP-2, while identifying PPEP-2’s endogenous substrate as the most optimal substrate in our library data. Furthermore, by employing the library approach, we investigated the altered specificity of mutants of PPEP-1 and PPEP-2.

Additionally, we characterized a novel PPEP from *Anoxybacillus tepidamans*, which we termed PPEP-4. Based on structural comparisons, we hypothesized that PPEP-4 displays a PPEP-1-like prime-side specificity, which was substantiated by the experimental data. Intriguingly, another putative PPEP from *Clostridioides difficile*, CD1597, did not display Pro-Pro endoproteolytic activity.

Collectively, we characterized PPEP specificity in detail using our robust peptide library method and, together with additional structural information, provide more insight into the intricate mechanisms that govern protease specificity.

## Introduction

Proteases represent a diverse and indispensable class of enzymes that play pivotal roles in cellular homeostasis, protein turnover, and the regulation of various biological pathways. Their ability to hydrolyze peptide bonds is vital for the activation, maturation, or degradation of proteins. Among the diverse array of proteases found across different organisms, bacterial proteases are particularly intriguing due to their significance in bacterial physiology [1,2], pathogenesis [3,4], antimicrobial drug targets [5] and biotechnological applications [6].

Protease function is largely determined by substrate specificity, i.e., which residues are tolerated surrounding the cleavage site. Proline residues, for instance, are generally excluded as part of the cleavage site due to their cyclic structure, imposing conformational constraints that hinder proteolytic cleavage [7,8]. However, several proteases have been described that selectively cleave N- or C-terminally of proline residues [9–12]. A notable group of bacterial proteases, the Pro-Pro endopeptidases (PPEPs), possess the unique ability to specifically hydrolyze proline-proline bonds.

PPEPs are predicted to be present in many bacterial species [13] and several PPEPs have been characterized [14–16]. Although these enzymes appear very similar based on their protein sequence, small structural differences result in distinct substrate specificities [16]. PPEP specificity is at the minimum dependent on the six residues flanking the cleavage site (P3-P2-P1↓P1’-P2’-P3’, Schechter and Berger nomenclature [17]), with the permissible residues dictated by interactions within the active site. A comprehensive understanding of PPEP specificity holds promise for predicting endogenous substrates, facilitating industrial applications, developing inhibitors, and their use as potential biomarkers.

Several methods are available to profile protease specificity, such as gel- and fluorescence-based methods [18], N-terminomics approaches [19,20], and library methods. For the latter, approaches using phage display [21,22], positional scanning [23], proteome-derived libraries [24] and synthetic combinatorial peptide libraries exist [25,26]. Synthetic combinatorial peptide libraries consist of systematically synthesized peptides that cover all possible amino acid combinations around a cleavage site. Compared to proteome-derived peptide libraries, synthetic combinatorial peptide libraries contain potential substrates in equimolar amounts, which allows for a more quantitative approach.

Previously, we reported a novel method to profile PPEP prime-side specificity by combining the use of a synthetic combinatorial peptide library with LC-MS/MS analysis [16]. In this approach, protease-generated product peptides are enriched by negative selection and subsequently analyzed by LC-MS/MS. This approach allowed for an in-depth characterization of the prime-side specificity of PPEP-1, PPEP-2, and PPEP-3 and revealed the differences between the three PPEPs. However, PPEP specificity is also determined by the substrate’s non-prime-side residues. To obtain a thorough understanding of PPEP specificity, a method that allows for the characterization of the non-prime-side specificity is needed as well.

In order to achieve an integrated analysis of both non-prime- and prime-side specificities, we expanded our combinatorial peptide library method by synthesizing a complementary library that allowed us to profile the non-prime-side specificity of PPEPs. In addition, profiling of the complete specificity of PPEPs was achieved by combining the non-prime- and prime-side libraries. We not only used our method with known PPEPs but also applied it to determine the specificity of two uncharacterized PPEP homologs from both *C. difficile* and *Anoxybacillus tepidamans*. By combining the specificity profiles of PPEPs with structural information, we elaborate on the structure-function relationship of PPEPs.

## Experimental procedures

### Expression and purification of recombinant PPEPs

PPEP-1, PPEP-2, PPEP-1_SERV_, and PPEP-2_GGST_ were expressed and purified as previously described [15,27]. For the expression of PPEP-4, a pET-16b vector containing an *E. coli* codon-optimized 10xHis- PPEP-4 (lacking the signal peptide) construct was ordered from GenScript. The pET-16b 10xHis-PPEP-4 plasmid was transformed to *E. coli* strain C43 and PPEP-4 expression was induced using 1 mM IPTG for 4 h. Bacterial pellets were resuspended in lysis buffer (20 mM NaH_2_PO_4_, 500 mM NaCl, 40 mM imidazole, pH 7.4) and lysozyme was added to the suspension to 1 mg/ml and incubated for 30 min on ice before disruption by 5 30 s rounds of sonication. The lysates were loaded onto a 1 ml HisTrap HP column (GE Healthcare) coupled to an ÄKTA Pure FPLC system (GE Healthcare). The column was washed using wash buffer (20 mM NaH_2_PO_4_, 500 mM NaCl, 40 mM imidazole, pH 7.4) and 10xHis-PPEP-4 was eluted using a linear gradient with elution buffer (20 mM NaH_2_PO_4_, 500 mM NaCl, 500 mM imidazole, pH 7.4). Buffer exchange to PBS was performed using an Amicon® Ultra-15 Centrifugal Filter Unit with a 10 kDa cut-off membrane.

For CD1597 (and the predicted catalytic domain, AA 211-416), a pET-28a vector containing an *E. coli* codon-optimized 6xHis-CD1597 (lacking the signal peptide) construct was ordered from Twist Bioscience. Expression and purification were performed as described above but with different lysis (50 mM NaH_2_PO_4_, 300 mM NaCl, 10 mM imidazole, pH 8.0), wash (50 mM NaH_2_PO_4_, 300 mM NaCl, 20 mM imidazole, pH 8.0), and elution (50 mM NaH_2_PO_4_, 300 mM NaCl, 250 mM imidazole, pH 8.0) buffers.

### Combinatorial peptide library assays

The combinatorial peptide libraries were synthesized and assays were performed as previously described [16]. In short, approximately 10 nmol of precleaned (on avidin column) peptide mixture was incubated with 200 ng PPEP for 3 h at 37 °C in PBS. For PPEP-1_SERV_ and PPEP-2_GGST_, 500 ng was used in combination with an incubation time of 16 h. A non-treated control was included. After incubation, the samples were loaded onto an in-house constructed column consisting of a 200 μL pipet tip containing a filter and a packed column of 100 μL of Pierce High Capacity Streptavidin Agarose beads (Thermo, the column was washed four times with 150 μL of PBS before use), to remove the biotinylated peptides. The flow-through and four additional washes with 125 μL PBS were collected. The product peptides were desalted using reversed-phase solid-phase extraction cartridges (Oasis HLB 1 cm3 30 mg, Waters) and eluted with 400 μL of 30% acetonitrile (v/v) in 0.1% formic acid. Samples were dried by vacuum concentration and stored at −20 °C until further use. For the peptide library assays in which the non-prime- and prime-side libraries were combined, approximately 5 nmol of each library was used (10 nmol in total).

### LC-MS/MS analyses

For the analyses of the product peptides of P3=Val non-prime-side sublibrary after incubation with PPEP-1 and those of the non-prime-side library after incubation with PPEP-1 and -2, product peptides were analyzed by online C18 nanoHPLC MS/MS with an Ultimate3000nano gradient HPLC system (Thermo, Bremen, Germany), and an Exploris480 mass spectrometer (Thermo). Peptides were injected onto a precolumn (300 μm × 5 mm, C18 PepMap, 5 μm, 100 A), and eluted via a homemade analytical nano-HPLC column (30 cm × 75 μm; Reprosil-Pur C18-AQ 1.9 μm, 120 A; Dr. Maisch, Ammerbuch, Germany). The gradient was run with a gradient of 2% to 36% solvent B (20/80/0.1 water/acetonitrile/formic acid (FA) v/v) in 52 min. The nano-HPLC column was drawn to a tip of ∼10 μm and acted as the electrospray needle of the MS source. The mass spectrometer was operated in data-dependent MS/MS mode for a cycle time of 3 s, with HCD collision energies at both 17V and 23V and recording of the MS2 spectrum in the Orbitrap, with a quadrupole isolation width of 1.2 m/z. In the master scan (MS1) the resolution was 120,000, the scan range 350–1600, at standard AGC target at a maximum fill time of 50 ms. A lock mass correction on the background ion m/z = 445.12003 was used. Precursors were dynamically excluded after n = 1 with an exclusion duration of 10 s and with a precursor range of 10 ppm. Charge states 1–5 were included. For MS2 the first mass was set to 110 Da, and the MS2 scan resolution was 30,000 at an AGC target of 100% @maximum fill time of 60 ms.

For the analyses of the product peptides of the mixed non-prime- and prime-side libraries following incubation with PPEPs (and separate analyses for CD1597), product peptides were analyzed by online C18 nano-HPLC MS/MS with a system consisting of an Easy nLC 1200 gradient HPLC system (Thermo, Bremen, Germany) and an Orbitrap Fusion LUMOS mass spectrometer (Thermo). Peptides were injected onto a homemade precolumn (100 μm × 15 mm; Reprosil-Pur C18-AQ 3 μm, Dr Maisch, Ammerbuch, Germany) and eluted via a homemade analytical nano-HPLC column (30 cm × 75 μm; Reprosil-Pur C18-AQ 1.9 μm). The gradient was run from 2% to 40% solvent B (20/80/0.1 water/acetonitrile/formic acid (FA) v/v) in 52 min. The nano-HPLC column was drawn to a tip of ∼5 μm and acted as the electrospray needle of the MS source. The LUMOS mass spectrometer was operated in data-dependent MS/MS mode for a cycle time of 3 s, with HCD collision energies at 20 V, 25V, and 30V and recording of the MS2 spectrum in the orbitrap, with a quadrupole isolation width of 1.2 m/z. In the master scan (MS1) the resolution was 120,000, the scan range 350–1600, at an AGC target of 400,000 at a maximum fill time of 50 ms. A lock mass correction on the background ion m/z = 445.12003 was used. Precursors were dynamically excluded after n = 1 with an exclusion duration of 10 s and with a precursor range of 10 ppm. Charge states 1–5 were included. For MS2 the first mass was set to 110 Da, and the MS2 scan resolution was 30,000 at an AGC target of 100% @maximum fill time of 60 ms.

### LC-MS/MS data analysis

To identify product peptides in a database search after analysis of the non-prime-side library with PPEP-1 and -2, we generated a database containing all 130,321 possible peptides in the library, i.e., PTEDAVXXPPXXE-Ahx-Ahx-K (biotin was included as a variable modification in the database searches). The Ahx in all peptide sequences was replaced by a Leu (they have an identical mass). For the identification of product peptides after analysis of the mixed non-prime- and prime-side libraries, a database was generated containing all 9-mer product peptides that are possible based on Pro-Pro cleavage (i.e., PTEDAVXXP and PXXGGLEEF).

Raw data were converted to peak lists using Proteome Discoverer version 2.5.0.400 (Thermo Electron) and submitted to the in-house created databases using Mascot v. 2.2.7 (www.matrixscience.com) for peptide identification, using the Fixed Value PSM Validator. Mascot searches were with 5 ppm and 0.02 Da deviation for precursor and fragment mass, respectively, and no enzyme specificity was selected. Biotin on the protein N-terminus was set as a variable modification.

The database search results were filtered for product peptides that contained either PTEDAV or GGLEEF, were 9 residues in length, and contained no biotin. The resulting peptide lists were transported to Microsoft Excel, where duplicate masses and corresponding abundances were removed (e.g., the abundances of isomers PLPGGLEEF and PIPGGLEEF are listed twice, while this abundance is the total abundance of the two variants). The most abundant product peptides that together accounted for >90% of the total abundance were selected for further analysis (except for PPEP-4; see Results and Discussion). Further analysis was performed in Skyline 23.1.0.268 by importing the product peptides as FASTA along with the raw data files [28]. The Extracted Ion Chromatograms (EICs) displaying the product peptides were created by plotting the intensities of the signals corresponding to the monoisotopic m/z values of both 1+ and 2+ charged peptides with a mass tolerance of 5 ppm. Peptide annotation in Skyline was refined by manual inspection of MS/MS spectra and peak areas were exported from Skyline and used to create the sequence logos using WegLogo 3.7.12 [29].

### FRET peptide cleavage assays

Time course kinetic experiments with PPEPs were performed using fluorescent FRET-quenched peptides. FRET peptides consisted of Lys(Dabcyl)-EXXPPXXD-Glu(EDANS), in which each X varied between the different peptides tested. Proteolysis of FRET peptides by PPEPs was tested in 150 µl PBS containing 50 mM FRET peptide and 200 ng enzyme. Peptide cleavage was measured using the Envision 2105 Multimode Plate Reader. Fluorescence intensity was measured each minute for 1 h, with 10 flashes per measurement. The excitation and emission wavelengths were 350 nm and 510 nm, respectively. For the assay at different pH, buffers were prepared by mixing 0.2 M NaH_2_PO_4_ and 0.2 M Na_2_HPO_4_ and adding dH_2_O to dilute the buffer 2x. The resulting buffers had a pH of 5.8, 6.4, 7.0, 7.5, and 8.0.

### Bioinformatic analyses

Signal peptide predictions were performed using SignalP 6.0 [30]. Sequence alignments were performed using the Clustal Omega Multiple Sequence Alignment tool [31]. The predicted structures of PPEP-3 and PPEP-4 were retrieved from the Alphafold DB (https://alphafold.ebi.ac.uk/). Structures were analyzed using PyMOL (The PyMOL Molecular Graphics System, Version 2.5.5 Schrödinger, LLC).

## Results and Discussion

### Design and testing of a synthetic combinatorial peptide library to determine PPEP non-prime-side specificity

To determine the non-prime-side specificity of PPEPs, we constructed a new synthetic combinatorial peptide library according to the sequence PTEDAVXX**PP**XXEZZO motif (X=any residue except Cys, Z=6- aminohexanoic acid, O=Lys(biotin)-amide) (**Figure 1A**). In analogy with the previous library that was used to determine the prime-side specificity of PPEPs (**Figure 1A**) [16], the two core Pro residues (P1- P1’) were fixed while surrounding positions (P3-P2, P2’-P3’) could contain any amino acid residue (except Cys, omitted to prevent disulfide bridges). In contrast to the prime-side library, the biotin in the new library was added at the C-terminus, while the peptide tail was added to the N-terminus. The sequence of this tail (PTEDAV) showed good chromatographic behavior and fragmentation characteristics (**Supplemental Figures S1 and S2**) and was based on product peptides from the endogenous substrates of PPEP-1 [27]. Collectively, the new library was designed to allow for the identification of PTEDAVXXP product peptides following a similar approach as previously described (**Figure 1B**) [16].

**Figure 1.**
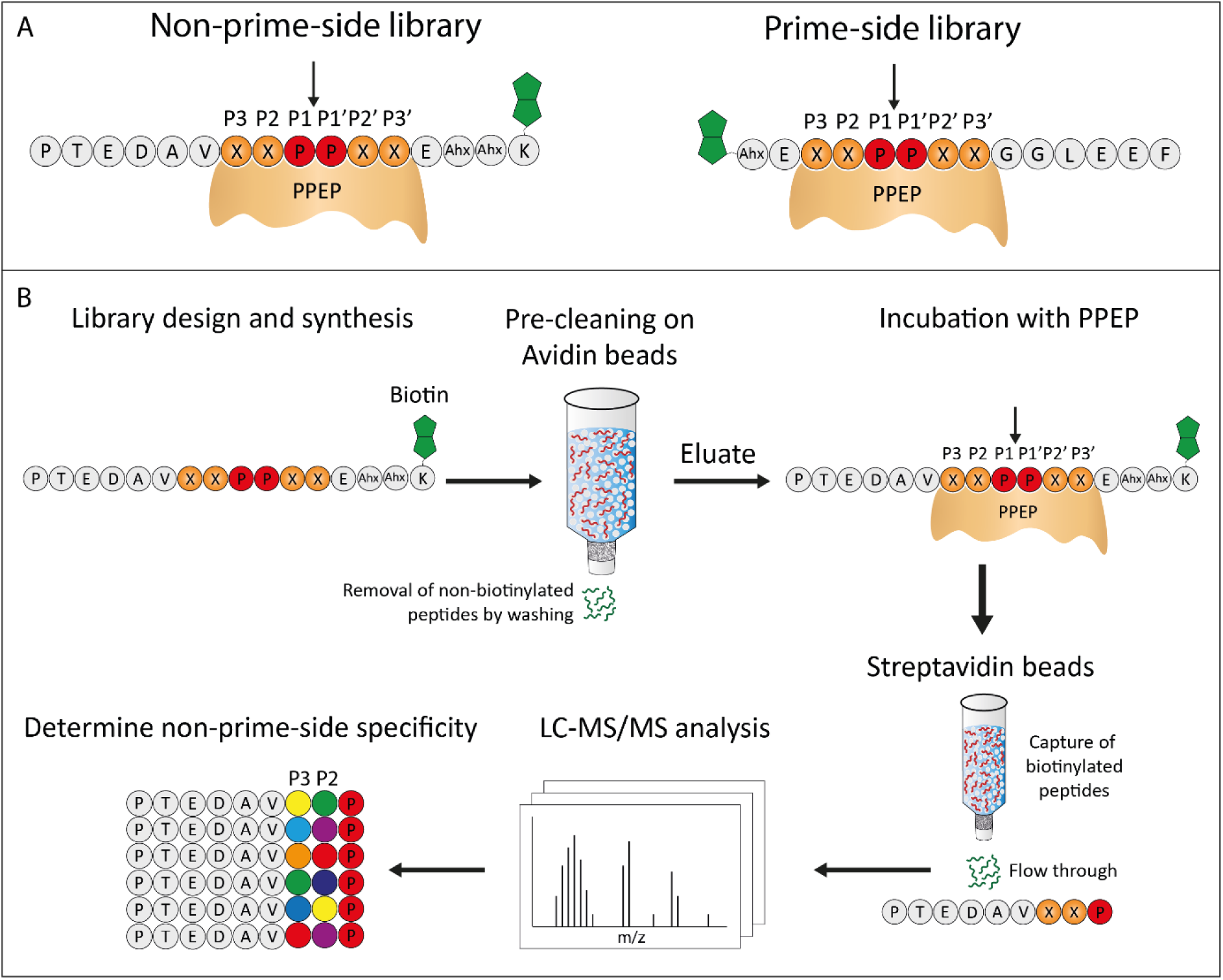
Design of the non-prime-side synthetic combinatorial peptide library. **A)** Design of the non-prime-side library (left) and the previously described prime-side library (right) [16]. The expected cleavage site is indicated with an arrow and biotin is represented in green. Ahx=6-aminohexanoic acid. **B)** Strategy for determining the non-prime-side specificity of PPEPs. Nonbiotinylated peptides are removed by washing the peptide library on an avidin column. Then, the eluted peptide library is incubated with a PPEP and subsequently loaded onto a streptavidin column. The biotinylated peptides are captured, while the PTEDAVXXP product peptides pass through the column. The product peptides are analyzed by LC-MS/MS, after which the non-prime-side specificity can be determined. Figure was adapted from Claushuis et al. (2023) [16], which is available under Creative Commons Attribution 4.0 International License.

To test the specificity profiling potential of the newly synthesized peptide library, we incubated the library with PPEP-1 and PPEP-2 (**Figure 2**). Previously, we reported that PPEP-1 and PPEP-2 have a markedly different non-prime-side specificity since the proteases are unable to cleave each other’s substrates [15]. In addition, PPEP-1 is known to tolerate multiple amino acids at the P2 and P3 positions [27], while for PPEP-2 the non-prime-side specificity has been less explored. The data in **Figure 2** corroborated the difference in non-prime specificity between PPEP-1 and PPEP-2, since most of the product peptides were not shared between the two PPEPs. In addition, it was readily apparent that the non-prime-side specificity was less stringent for PPEP-1 than for PPEP-2, i.e., more different product peptides were needed to account for >90% of the total intensity.

**Figure 2.**
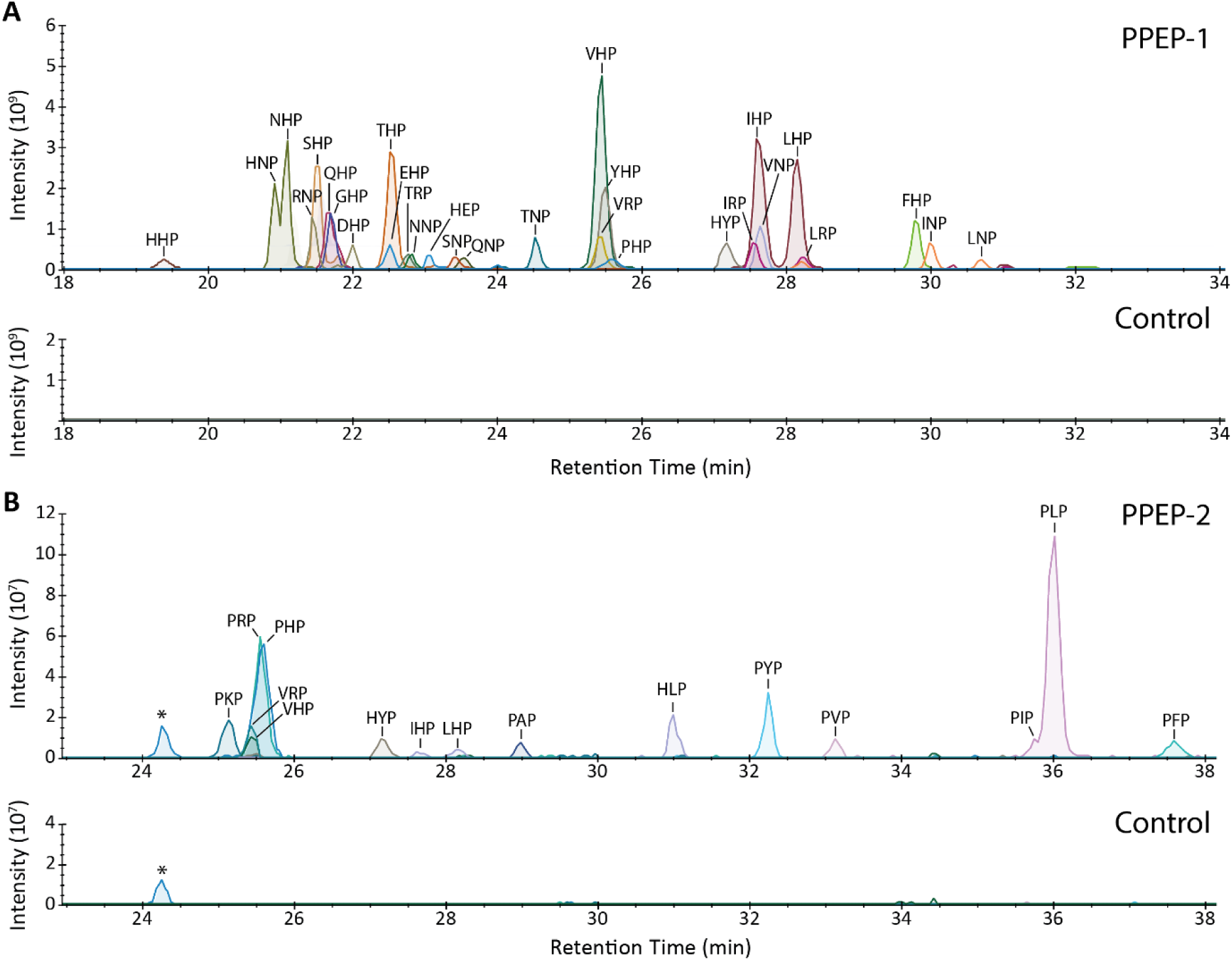
Incubation of the non-prime side combinatorial library with PPEP-1 and PPEP-2. Peptides were incubated with either **A)** PPEP-1 or **B)** PPEP-2. After product peptide enrichment and analysis with LC-MS/MS, a database search was performed using an in-house created database containing the 130.321 possible 16-mer peptides. The results were filtered for the 9-mer product peptides corresponding to the cleavage between the two fixed prolines (PTEDAVXXP). From these, the most abundant product peptides that together accounted for >90% of the total intensity were used to create an EIC. To distinguish between isomeric product peptides, manual inspection of MS/MS spectra, combined with LC-MS/MS of additional synthetic peptides, was performed. Mass tolerance was set to 5 ppm. An untreated control sample was included. *A molecule corresponding to the mass of PTEDAVPHP (481.7325, [M+2H]^2+^) was observed but MS/MS spectra indicate no PTEDAVXXP product peptide.

Also with the new library, the assignment of specific product peptides was based on manual inspection of MS/MS spectra and additional LC-MS/MS analyses of candidate product peptides. For example, this allowed us to unambiguously assign the major product peptide of PPEP-2 as PTEDAVPLP and not PTEDAVPIP (**Figure 2** and **Supplemental Figure 3A**). Moreover, a cleavage assay with PPEP-2 and FRET-quenched peptides showed that PLPPVP is cleaved much more efficiently than PIPPVP by PPEP-2 (**Supplemental Figure S4A**). Similar analyses were used to correctly assign other isomeric peptides, e.g. PTEDAVHIP, PTEDAVHLP, PTEDAVIHP and PTEDAVLHP (**Supplemental Figure S3B and S4B**).

We also removed two 9-mer product peptides from our analyses, PTEDAVGGP and PTEDAVAGP, because manual inspection of the data demonstrated that they actually correspond to the isomeric 8-mer peptides PTEDAVNP and PTEDAVQP, resulting from cleavage before the first fixed Pro at P1 in our design (PTEDAVXX↓PPXX).

Although the product peptide signals in the PPEP-2 treated sample and the untreated control greatly differ due to the proteolysis by PPEP-2, the signals in the PPEP-2 treated sample are approximately two orders of magnitude lower compared to the signals we observe in the PPEP-1 treated sample (**Figure 2**). Previously, we observed roughly five times lower signals for PPEP-2 prime-side product peptides compared to those of PPEP-1 [16]. This is a markedly smaller difference than we have observed in our non-prime-side results. A likely explanation is that, either for PPEP-1 or PPEP-2, the P4 or the P4’ position also determines substrate specificity, since these differ in the designs of both peptide libraries while all other factors remained constant (**Figure 1**).

Overall, the results with PPEP-1 and PPEP-2 as described above were in line with our expectations and showed that profiling of PPEP specificity could be achieved with the non-prime-side peptide library.

### Profiling the non-prime- and prime-side specificity of PPEPs in a single experiment

Having tested the new library, we next sought to profile the P3-P3’ specificity of PPEPs in a single experiment. To this end, we mixed the previously described prime-side peptide library [16] with the newly synthesized non-prime-side library (1:1) and incubated the mixture with either PPEP-1 or PPEP-2. The total amount of peptides was left unaltered, meaning that in comparison to the non-prime-side library experiment, only half the amount of each peptide was used. The results of the LC-MS/MS analyses were used to create EICs of the product peptides (**Figure 3A,B**). Based on the intensities of the product peptides, we constructed logos depicting the relative occurrence of a residue at a position surrounding the cleavage site (**Figure 3C,D**). The logos show how strongly the specificity at a position surrounding the cleavage site is determined by certain residues. Although the logos in **Figures 3C,D** do not take subsite cooperativity into account (they only show the relative occurrence of a residue at a certain position in the product peptides), this is the case for the EICs (**Figure 3A,B**). Therefore, the EICs and logos are complementary to each other. Since we make use of two different peptide libraries, we cannot draw conclusions about the subsite cooperativity spanning the cleavage site, e.g., the influence of a residue at the P2 position on the tolerance by a protease for a residue at the P2’ position.

**Figure 3.**
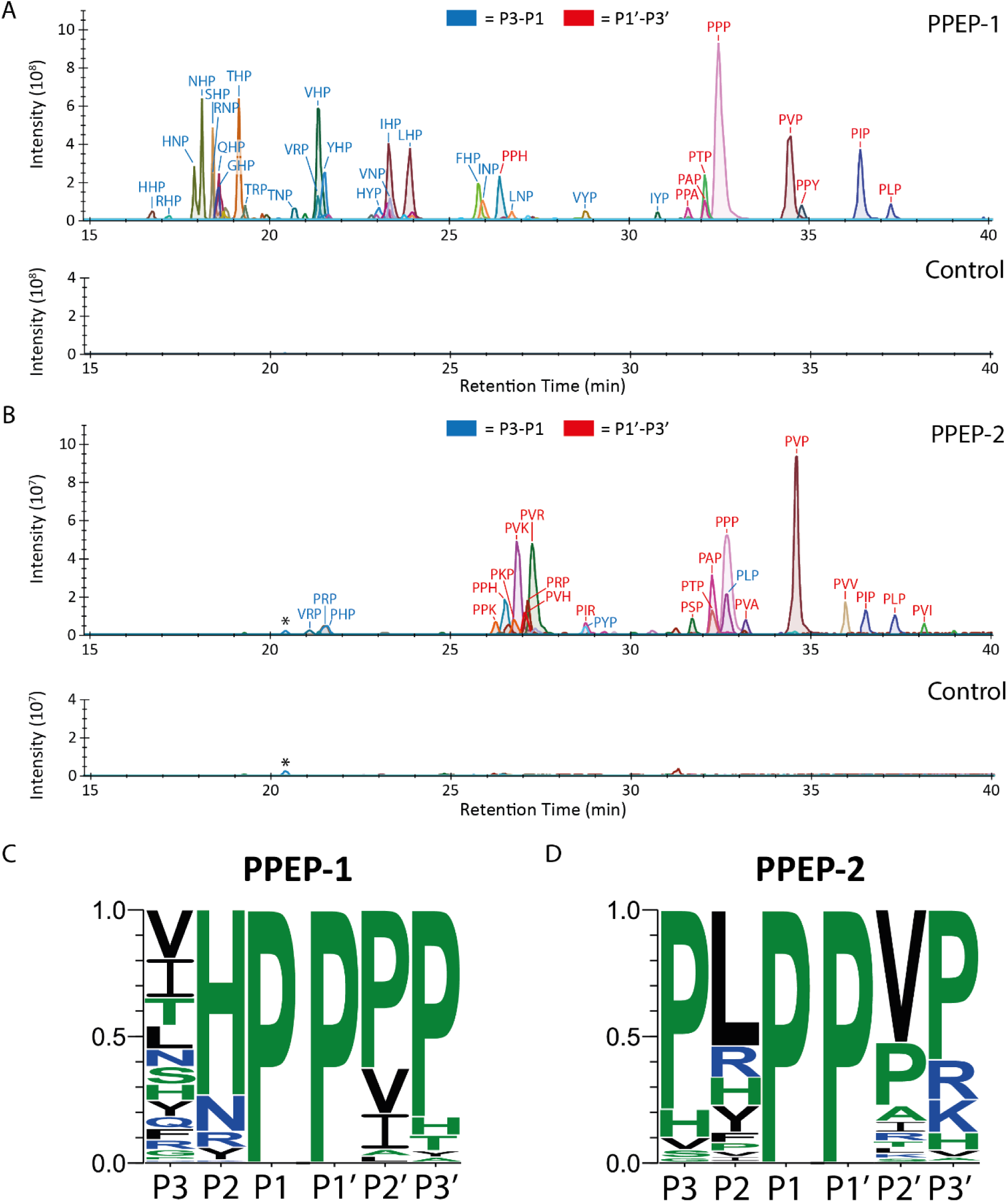
Incubation of the mixed non-prime- and prime-side libraries with PPEP-1 and PPEP-2. The non-prime- and prime-side libraries were mixed and the peptides were incubated with either **A)** PPEP-1 or **B)** PPEP-2. The product peptides were analyzed using LC-MS/MS. Results were searched against a database containing all 722 9-mer product peptides based on cleavage between the two fixed prolines, i.e., PTEDAVXXP and PXXGGLEEF (X≠Cys). The most abundant products that together account for >90% of the total abundance per library were used to create the EICs. Mass tolerance was set to 5 ppm. An untreated control sample was included. **C) and D)** The results from the EICs were used to create a logo that displays the observed frequency of a residue at positions P3-P3’ for PPEP-1 **(C)** and PPEP-2 **(D)**. *A molecule corresponding to the mass of PTEDAVPHP (481.7325, [M+2H]^2+^) was observed but MS/MS spectra indicate no PTEDAVXXP product peptide.

First of all, the new data for the prime-side specificity of PPEP-1 and PPEP-2 are consistent with our previously reported data [16], thereby demonstrating the excellent reproducibility of the method, even with the inclusion of the new combinatorial peptide library in the experiment (**Figure 3C and D)**.

Notably, for PPEP-1, the logo highlights the variability of permissible residues at the P3 position. This observation aligns with biological expectations, since the endogenous substrates (CD2831 and CD3246) have a Val, Ile, or Leu at the P3 position, suggesting a lower stringency at this position for PPEP-1 activity in *C. difficile* [14]. In the context of an Asn at the P2 position, as is the case in the endogenous substrates of PPEP-1, product peptides containing TNP and HNP (although the intensity of this signal is influenced by the ESI response factor [16,32]) are among the highest signals aside from VNP (**Figure 2A and 3A**). In the PPEP-1 cocrystal with substrate VNPPVP (P3-P3’), the Tyr94, Leu95, Trp110, and Leu116 are found in close proximity to the Val (P3) and likely influence the P3 specificity (**Supplemental Figure 5A**) [33]. Substitution of the Val (P3) with a Thr in the PPEP-1 cocrystal had no consequences, because of their similar sizes and the absence of attractive or repulsive polar interactions (**Supplemental Figure 5B**). However, substituting the Val (P3) with His could produce polar interactions with the Tyr94 in PPEP-1 (**Supplemental Figure 5C**), potentially strengthening the interaction between protease and substrate. In this configuration, the His side chain extended away from the P3 contacting residues, which might be a common mechanism to mitigate steric clashes and could explain the high variability of residues at the P3 position.

In line with the data presented above **(Figure 2)**, the PPEP-1 logo shows a high preference for His at the P2 position (**Figure 3C**). This is surprising since this residue is not observed at the P2 position in the endogenous substrates. The high abundance of His and the other basic amino acids (Arg and Lys) in the logos could relate to their ionization efficiency [32]. To assess how well His is tolerated at the P2 position compared to Asn and the other basic amino acids, a cleavage assay using FRET-quenched peptides was performed (**Supplemental Figure S6**). Although the His at the P2 position is well tolerated by PPEP-1, an Asn at that position produces the optimal substrate. In line with **Figure 3A**, VRP (P3-P1) is cleaved less efficiently than VHP, while VKP represents a very poor substrate.

The preference of PPEP-1 for Asn at the P2 stems from the interactions of the side chains of Lys101 with the side chains of Glu184, Glu185, and the Asn at the P2 position, collectively termed the KEEN interface [33]. Modeling of a PPEP-1 cocrystal with a substrate in which the Asn has been substituted for a His residue reveals that the histidine side chain can interact with Lys101 via hydrogen bonding as well (**Figure 4A**). In addition, the backbone atoms of the His interact with Gly117, similar to Asn. In the interaction as shown in **Figure 4A**, the electronegative nitrogen (N1) of His interacts with the protonated Lys101. However, in case of His protonation, this interaction might be lost. To test this, we incubated PPEP-1 with the substrate VHPPPP at various pH values (**Figure 4B**). At pH 5.8, we expected most of the His to be protonated since the pKa of the His side chain is 6.04. To see the effect of His protonation rather than the effect of a lower pH on the PPEP activity in general, we also included the substrate VNPPPP as a control. When comparing the initial slope of the reactions, the reactions of both substrates were similar at pH 7.5 and 8.0. However, by lowering the pH, VHPPPP cleavage became increasingly less efficient compared to VNPPPP, and at pH 5.8, the initial slope of the VHPPPP reaction was around 6x lower than that of VNPPPP. This indicated that protonation of the His at P2 inhibits cleavage by PPEP-1, likely due to the loss of the interaction with the Lys101.

**Figure 4.**
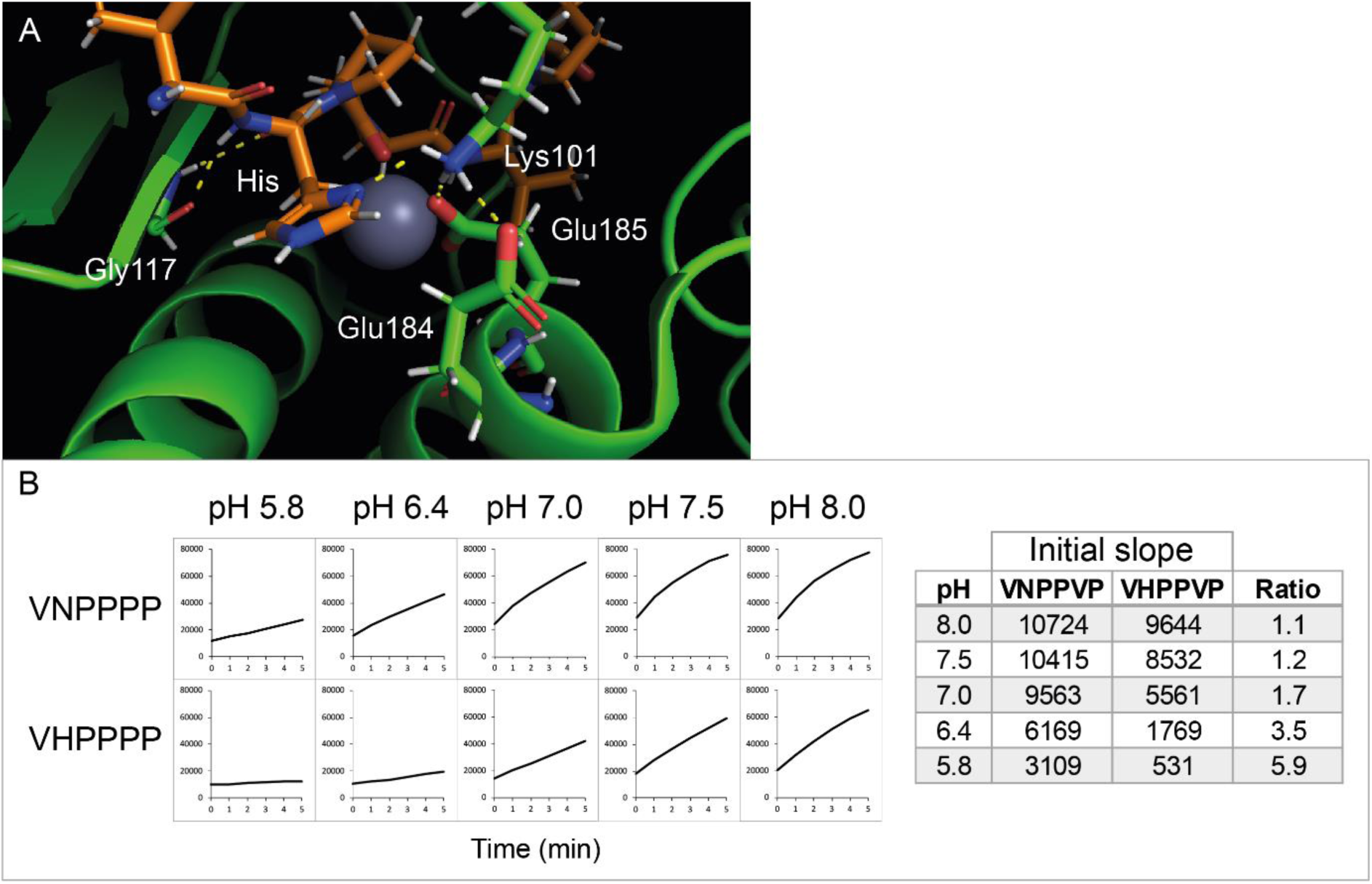
Tolerance for His at the P2 position by PPEP-1 is dependent on the pH. **A)** Cocrystal structure of PPEP-1 (green, PDB: 5A0X) with a substrate peptide (orange) in which the Asn is substituted by a His. Yellow dotted lines indicate the interactions between residues. **B)** Time course of the PPEP-1 mediated cleavage at different pH with FRET-quenched peptides Lys(Dabcyl)- EV(H/N)PPPPD-Glu(EDANS) (left). The initial slope represents the increase in fluorescence in the first 5 min. The ratios of the initial slopes were calculated to compare both reactions (right).

Importantly, the results of our synthetic combinatorial peptide library approach demonstrate the ability to identify endogenous substrates using this method as **Figures 3B,D** clearly show the preference of PPEP-2 for PLPPVP (P3-P3’). When forming a hypothesis about the endogenous substrates without any other prior knowledge, a search for this motif in the proteome of *P. alvei* directly leads to the identification of the endogenous substrate VMSP of PPEP-2 [14]. Previous modeling of PPEP-2 with the endogenous substrate PLP↓PVP (P3-P3’) predicted the Pro at the P3 to produce a kink in the polypeptide, thereby redirecting the upstream polypeptide away from the salt bridge formed by Glu113 and Arg145 [15]. The need for this diversion was supported by the data in **Figure 3B**, since the presence of a Pro at the P3 position was a strong determinant for proteolytic activity.

### Exchanging the β3/β4 loop of PPEP-1 and PPEP-2 shifts specificity towards one another

While PPEP-1 and PPEP-2 share a close relationship, their non-prime-side specificity exhibits marked differences. Among other structural elements, the β3/β4 loop plays a role in non-prime-side specificity [15]. Notably, this loop varies largely between the two proteases. Previous studies demonstrated that replacing the β3/β4 loop of PPEP-2 (^112^SERV^115^) with that of PPEP-1 (^117^GGST^120^) alters the enzyme’s preference for the P3 position, shifting from Pro to Val and making it more PPEP-1-like [15].

Surprisingly, this effect was not mirrored in PPEP-1, as the mutant PPEP-1_SERV_ failed to cleave the tested peptides VNPPVP and PLPPVP. To gain a more comprehensive understanding of the significance of the β3/β4 loop for PPEP specificity, we conducted experiments using the combined non-prime- and prime side libraries incubated with PPEP mutants (PPEP-1_SERV_ and PPEP-2_GGST_). EICs of the non-prime-side product peptides with VNP (PPEP-1 substrate), PLP (PPEP-2 substrate), and their combinations PNP and VLP, were generated (**Figure 5**).

**Figure 5.**
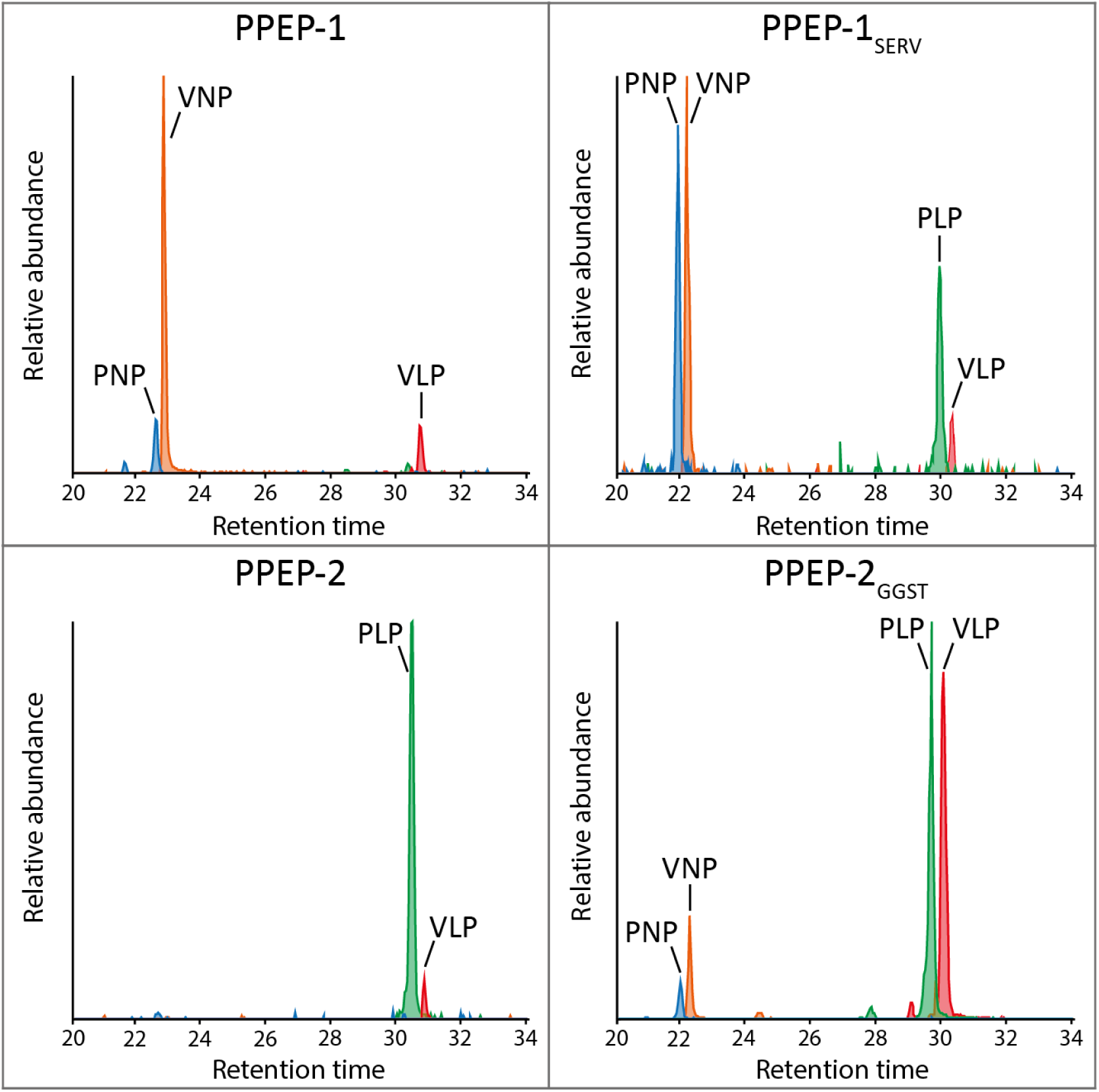
Altered non-prime-side specificity of PPEP-1_SERV_ and PPEP-2_GGST_. EICs were constructed for the product peptides containing the non-prime-side P3-P1 sequences VNP, PLP, VLP, and PNP after incubation of the mixed library with PPEP-1 (upper left panel), PPEP-2 (bottom left panel), PPEP-1_SERV_ (upper right panel), or PPEP-2_GGST_ (bottom right panel).

As expected, for wild-type (WT) PPEP-1, VNP was the predominant product peptide. However, the mutant PPEP-1_SERV_ displayed a shift in specificity towards PPEP-2, evidenced by increased signals for PNP and PLP product peptides, whereas the specificity for VLP remained unchanged. Conversely, PPEP-2_GGST_ exhibited a similar shift towards WT PPEP-1 specificity, with a relative increase in signals for PNP, VNP, and VLP compared to PPEP-2. Notably, the original substrates continued to be favored, suggesting that the β3/β4 loop is not the primary determinant of non-prime-side specificity. Instead, residues Lys101 and Glu184 in WT PPEP-1 that are part of the KEEN interface [33] and interact with the Asn at the P2 position (aligning with residues Arg96 and T180 in PPEP-2), may play a more decisive role in P3 and P2 specificity. Additionally, the β3/β4 loop in PPEP-2 is stabilized by a salt bridge between residues Glu113 and Arg145 [15]. However, this stabilizing interaction might be absent in PPEP-1_SERV_ due to the substitution of Arg145 with His in WT PPEP-1. Consequently, the steric hindrance caused by this salt bridge might be absent in PPEP-1_SERV_, potentially allowing a Val at the P3 position.

It’s worth noting that PPEP-1_SERV_, previously incapable of cleaving FRET-quenched peptides containing the VNPPVP and PLPPVP sequences [15], now demonstrated tolerance for both VNP and PLP at the non-prime-side. Prime-side specificity remained unaffected by the mutation, with PVP (P1’-P3’) product peptides present post-incubation with PPEP-1_SERV_ (**Supplemental Table S1**). Discrepancies between current and previous results may be attributed to differences in the P4 position, where the FRET-quenched peptides featured a Glu, while the non-prime-side library had a Val at that position.

### A second putative PPEP from C. difficile does not exhibit Pro-Pro endopeptidase activity

In *C. difficile*, a second PPEP-like protein, CD1597 (Gene: CD630_15970, UniProt ID: Q186F3), is present. Interestingly, CD1597 differs from the PPEPs that have been hitherto been described in several ways. Apart from a PPEP-like domain, CD1597 contains an additional N-terminal domain that makes up about half of the protein (**Supplemental Figure S7**). Moreover, in contrast to the other PPEPs, CD1597 is not predicted to contain a signal peptide for secretion. Furthermore, several amino acid insertions in CD1597 are observed in a sequence alignment with PPEP-1 and PPEP-2 (**Supplemental Figure S7**).

To assess the capability of CD1597 to hydrolyze Pro-Pro substrates, we conducted separate incubations of non-prime- and prime-side libraries with CD1597. Subsequent database searches aimed at identifying product peptides did not reveal the formation of any products. Therefore, to visualize the data, we constructed EICs for all the possible 9-mer product peptides (PTEDAVXXP and PXXGGLEEF) (**Supplemental Figure S8**). In contrast to the previous experiments, the intensity of signals in the treated samples was comparable to those in the control samples, indicating a lack of proteolytic activity of CD1597. In some cases, however, a signal was exclusively observed in the treated sample, but manual inspection of the MS/MS spectra indicated no product peptides of any kind. To rule out the possibility of an inhibitory effect of the N-terminal domain that is absent in other PPEPs, experiments with only the predicted proteolytic domain were conducted but yielded similar results (data not shown). Additional investigations into CD1597, using both FRET-quenched and plain peptide cleavage assays, consistently revealed no cleavage (data not shown). In light of all these findings, we conclude that, despite its structural resemblance to a PPEP, CD1597 does not exhibit PPEP-like activity.

The underlying cause of CD1597’s inactivity remains elusive. Although we cannot rule out the possibility that we have purified an inactive recombinant protease, we find this unlikely given the previous work on other PPEPs for which we use similar purification routines. While the absence of activity could suggest CD1597 is a pseudoprotease, it’s noteworthy that the protein harbors an intact catalytic HEXXH motif, which contradicts the typical characteristics of a pseudoprotease [34,35]. Interestingly, we have only identified a single peptide of CD1597 in two extensive proteome analyses of overnight cultures of *C. difficile* [36], proving that CD1597 has a very low abundance. However, CD1597 has been identified in larger quantities in the spore coat/exosporium layer of *C. difficile* spores [37]. In the context of *C. difficile* spores, two other pseudoproteases, namely CspA and CspB, have been identified, and they play crucial roles in spore germination [38]. The seemingly exclusive presence of CD1597 in spores could indicate a potential involvement in spore germination, either as a pseudoprotease or as a zymogen that requires specific stimuli for activation. Further exploration is needed to unravel the precise role of CD1597 in the context of *C. difficile* spores and to elucidate the factors influencing its apparent inactivity in our library assays.

### Specificity profiling of a novel PPEP from Anoxybacillus tepidamans

Bioinformatic analysis predicted the presence of a PPEP homolog in the thermophilic bacterium *Anoxybacillus tepidamans* (gene: HNQ34_002771, UniProt ID: A0A7W8IRZ3) [13]. This protein, hereafter designated as PPEP-4, showed a close phylogenetic relationship to the previously described PPEP-3 from *Geobacillus thermodenitrificans* [13]. In fact, *A. tepidamans* was initially proposed to be a member of the genus *Geobacillus*, further demonstrating the close relationship of these organisms [39].

Prior investigations into the prime-side specificity of PPEP-3 indicated a strong preference for prolines at the P2’ and P3’ positions, in contrast to the more versatile P2’ specificity of PPEP-1 and -2 (**Figure 3** and [16]). The primary structure of PPEP-4 very closely resembles that of PPEP-3, although some differences are observed (**Figure 6A**). Notably, the substitution of Phe190 in PPEP-3 with Leu in PPEP-4, similar to PPEP-1, stood out. In PPEP-1, this residue is close to the P2’ Val in the substrate VNP↓PVP [33]. Likely, the Phe190 in PPEP-3 causes steric hindrance for a Val and most other residues at the P2’ position (**Figure 6B**) and causes the need for a Pro to redirect the substrate elsewhere.

**Figure 6.**
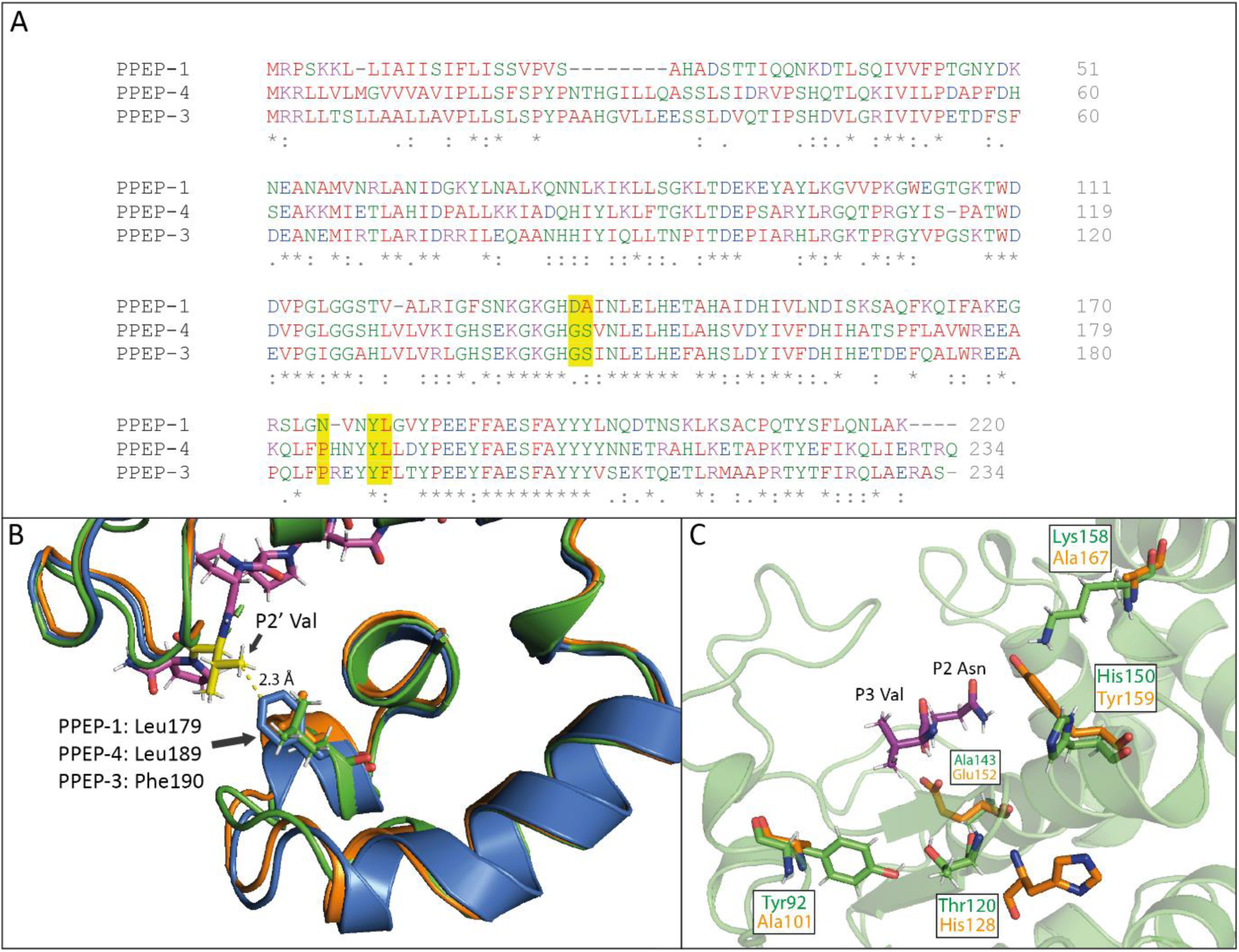
Comparison of PPEP-4 from *Anoxybacillus tepidamans* with PPEP-1 and PPEP-3. **A)** Sequence alignment of PPEP-1, PPEP-3, and PPEP-4 created using the Clustal Omega multiple sequence alignment tool. The residues in PPEP-1, and their corresponding residues in PPEP-3 and -4, that contact the Val at the P2’ in the substrate peptide VNPPVP are highlighted in yellow. **B)** Structural comparison of co-crystal of PPEP-1 (green, PDB: 6R5C) with its substrate VNPPVP (magenta, Val=yellow) and the predicted structures of PPEP-3 (blue) and PPEP-4 (orange). **C)** Comparison of the residues within 5 Å of the P3 Val and P2 Asn that differ biochemically between PPEP-1 (green) and the predicted structure of PPEP-4 (yellow).

We hypothesized that the substitution of Phe190 in PPEP-3 to Leu in PPEP-4 could render the P2’ specificity of PPEP-4 more akin to that of PPEP-1. To test this, we profiled PPEP-4 specificity using the combinatorial peptide libraries. Following LC-MS/MS analysis, a database search was performed using the database containing all possible 9-mer product peptides (PTEDAVXXP and PXXGGLEEF). In the experiments described above, we only included the most abundant product peptides that together accounted for >90% of the total intensity. However, in the case of PPEP-4, it became apparent that the non-prime-side specificity was highly diverse, and that the >90% cut-off value included too many substrates for meaningful visualization while many of them were very minor substrates. For clarity, we therefore decided to include only product peptides that represented at least 1% of the total intensity for both the non-prime- and prime-side product peptides. Using these product peptides, an EIC and logo were created to depict the P3-P3’ specificity of PPEP-4 (**Figure 7**).

**Figure 7.**
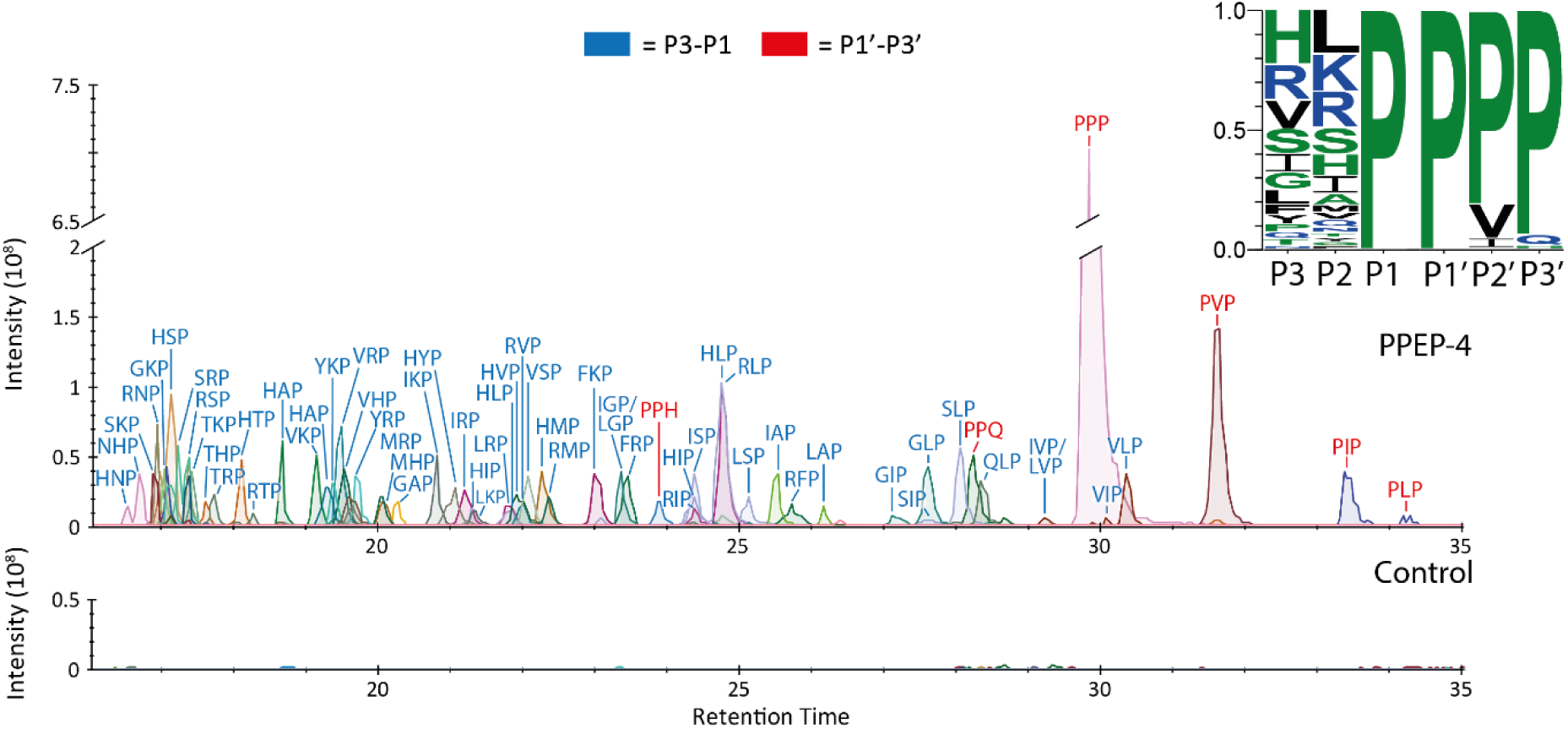
P3-P3’ specificity of PPEP-4 from *Anoxybacillus tepidamans*. The non-prime- and prime-side libraries were mixed and the peptides were incubated with PPEP-4. The product peptides were analyzed using LC-MS/MS and a database search was performed to identify and quantify the products. Results were filtered for 9-mer product peptides and the products accounting for >1% of the total abundance were used to create the EICs. Mass tolerance was set to 5 ppm. An untreated control sample was included. The results from the EICs were used to create a logo that displays the observed frequency of a residue at positions P3-P3’ for PPEP-4.

For the prime-side specificity of PPEP-4, PPP (P1’-P3’) is by far the most abundant product peptide, but also the PVP, PIP, and PLP product peptides are formed (**Figure 7**), resembling a PPEP-1-like specificity profile [16]. Since the substitution of the Phe190 in PPEP-3 to the Leu in PPEP-4 is the only difference between the P2’ contacting residues, we find it likely that the residue at this position is a large determinant for P2’ specificity. Furthermore, a product peptide containing the PPQ sequence (P1’-P3’) was found for PPEP-4, characteristic of PPEP-3 but not PPEP-1 [16]. Lastly, PPH (P1’-P3’) is tolerated by PPEP-4, similar to both PPEP-1 and PPEP-3. Initially, MS/MS spectra were inconclusive on the identity of the product peptide, i.e., it was either PPH or PHP. However, incubation of PPEP-4 with the FRET-quenched peptides containing VNPPPH and VNPPHP demonstrated that PPEP-4 exclusively cleaves VNPPPH, similar to PPEP-1 (**Supplemental Figure S9A**) [16]. Collectively, although PPEP-4 is highly similar to PPEP-3, PPEP-4’s prime-side specificity shows characteristics of both PPEP-1 and PPEP-3.

Although PPEP-1 tolerates many residues at the P3 and P2 positions (**Figures 3A,C**), PPEP-4 displays even greater flexibility (**Figure 7**). A comparison of the residues that are proximal to the P3 and P2 position in the PPEP-1 co-crystal with substrate VNPPVP and that differ biochemically between PPEP-1 and PPEP-4 (Alphafold model) revealed several residues that can influence P3-P2 specificity (**Figure 6C**). First, Tyr92 in PPEP is substituted by an Ala in PPEP-4. In addition, Thr120 is substituted by a His that is predicted to be further removed from the active site. Together, these differences might reduce the steric hindrances at the P3 (and possibly P4) positions. Furthermore, several other residues in PPEP-1 that are close to the P3-P2 residues are substituted in PPEP-4. Although these substitutions do not seem to reduce steric hindrances, their different biochemical properties might influence the intra- and intermolecular interactions that make the non-prime-side specificity of PPEP-4 more permissible. Further investigations of the structure-function relationships in PPEPs necessitate additional co-crystal structures.

A search for candidate substrates in the proteome of *A. tepidamans* focused on the motifs P↓PPP, P↓PVP, P↓PIP, P↓PPH, and P↓PPQ (↓=supposed cleavage site), resulting in the identification of 14 candidate substrates. Given the predicted secretion of PPEP-4, an analysis of these 14 candidates using SignalP 6.0 identified a single secreted protein, a predicted lipoprotein (gene: HNQ34_001056, UniProt ID: A0A7W8IQP4). This lipoprotein is predicted with high confidence to possess a Sec/SPII signal sequence for integration in the lipid membrane and contains its putative cleavage site (QNP↓PPP) close to the location of lipid insertion (**Supplemental Figure S10**). Although a FRET-quenched peptide with the core sequence QNPPPP is cleaved, proteolysis was incomplete after 1 h of incubation (**Supplemental Figure S9B**). A comparison with other peptides in our collection that were selected based on their high abundance at the non-prime-sides in the logo in **Figure 7** shows that QNP (P3-P1) is indeed poorly tolerated (**Supplemental Figure S9C**). However, the endogenous substrate of PPEP-4 does not necessarily need to possess the most optimal cleavage sequence, as is the case for PPEP-1 [27].

PPEP-1 and PPEP-2 are involved in adhesion/motility by cleaving large adhesive surface proteins, thereby releasing the cells. The lipoprotein in *A. tepidamans* differs from these substrates due to its small size and the lack of any predicted domains (or any other structural elements aside from the signal peptide). Another surface protein that does possess adhesion domains in the same organism and that contain PPEP-like cleavage motifs is the Penicillin-binding protein 1A (HNQ34_000435, UniProt ID: A0A7W8IMR6). The Penicillin-binding protein 1A is predicted to possess a transpeptidase domain, but also a Fibronectin type III (FN3) domain, which is known to be capable of binding components of the extracellular matrix, integrins, and possibly carbohydrates [40,41]. Interestingly, this protein contains a putative PPEP cleavage site directly upstream of the FN3 domain (EQPPAP) and two putative cleavage sites downstream of the FN3 domain (PTPPAP and TNPPAP), although these sites seem to represent poor substrates under our experimental conditions. Additional experiments such as bacterial surface-shaving [42] of *A. tepidamans* with PPEP-4 could provide more insight into the endogenous substrates of this member of the PPEP family.

## Conclusion

We developed an approach to characterize both the non-prime- and prime-side-specificity of PPEPs by combining the use of synthetic combinatorial peptide libraries with LC-MS/MS. Using this method, we deepened our understanding of the specificity of the previously characterized PPEP-1 and PPEP-2. Importantly, we were able to identify PPEP-2’s endogenous substrate sequence PLPPVP as the optimal substrate using our library method. In addition, we profiled the specificity of a novel PPEP from *A. tepidamans*, which we termed PPEP-4. Based on structural comparisons of PPEP-4 with other PPEPs, we predicted a P2’ specificity that resembles that of PPEP-1, which was confirmed by our data. Moreover, investigation of mutants of PPEP-1 and PPEP-2 that had their β3/β4 loop swapped showed that the non-prime-side specificity shifted towards each other, demonstrating the involvement of this loop in determining substrate specificity. For a second putative PPEP from *C. difficile*, however, no Pro-Pro endoproteolytic activity was observed. Finally, after including the non-prime-side peptide library, the prime-side profiles of PPEP-1 and -2 were in line with previously reported data, thereby demonstrating that the synthetic combinatorial library approach is a robust method with excellent reproducibility.

## Author contributions

P.J.H. and J.W.D. conceived the project. B.C. and P.J.H. performed experiments. B.C. and P.J.H. analyzed data. A.H.d.R., and J.v.A performed mass spectrometry analyses. P.A.v.V. provided the means for mass spectrometry analyses. B.C., P.J.H., J.W.D., and R.A.C. designed the library. R.A.C. produced the library. B.C. and J.C. performed protein expression and purification. B.C. and U.B. performed structural analyses and substrate modeling. P.J.H., M.W. and P.A.v.V supervised the project. P.J.H. and P.A.v.V acquired funding. B.C. visualized the results. B.C. and P.J.H. wrote the original draft. All authors reviewed and edited the paper.

## Acknowledgements

This research was supported by an ENW-M Grant (OCENW.KLEIN.103) from the Dutch Research Council (NWO) and by the research program Investment Grant NWO Medium with project number 91116004, which is (partially) financed by ZonMw. We thank Oleg I. Klychnikov and Stephen D. Weeks for the cloning and purification of PPEP-2, PPEP-1_SERV_, and PPEP-2_GGST_.

