## Supplemental Figures S1-S10 for "Non-prime- and Prime-side Profiling of Pro-Pro Endopeptidase Specificity Using Synthetic Combinatorial Peptide Libraries and Mass Spectrometry"

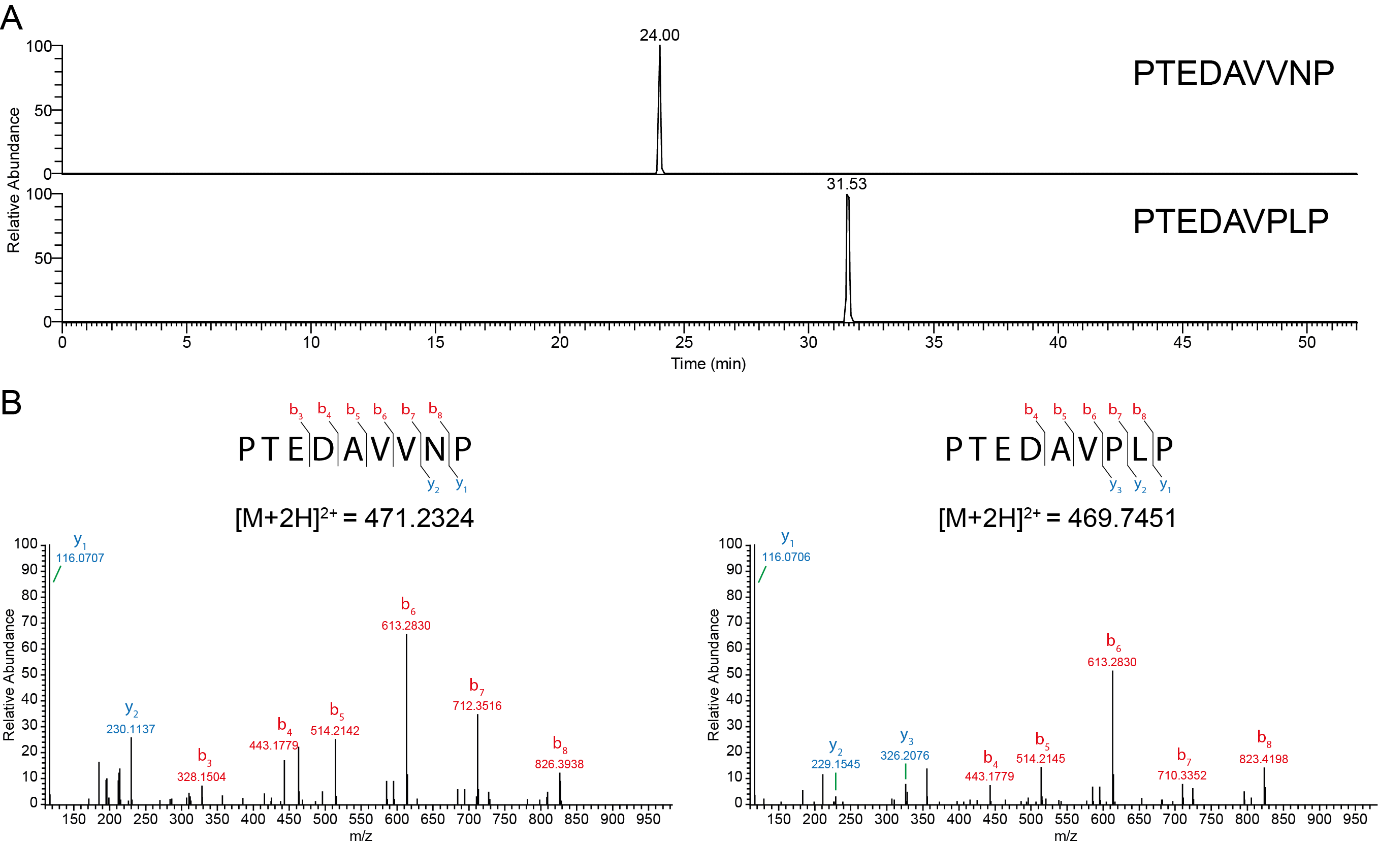

Figure S1. Characterization of the peptide tail for the library design**.** Peptides with the peptide tail PTEDAV and P3-P1 residues from PPEP-1 (VNP) and PPEP-2 (PLP) substrates were synthesized and analyzed by LC-MS/MS. **A)** Extracted Ion Chromatogram (EIC) of the peptides on a C18 column. The signals for the monoisotopic mass of the 1+ and 2+ charged species are shown with a mass tolerance of 10 ppm. **B)** MS/MS spectra of doubly charged species at an NCE of 23%. The b_7_ and y_2_ ions necessary for the correct assignment of the peptides were present in both MS/MS spectra.

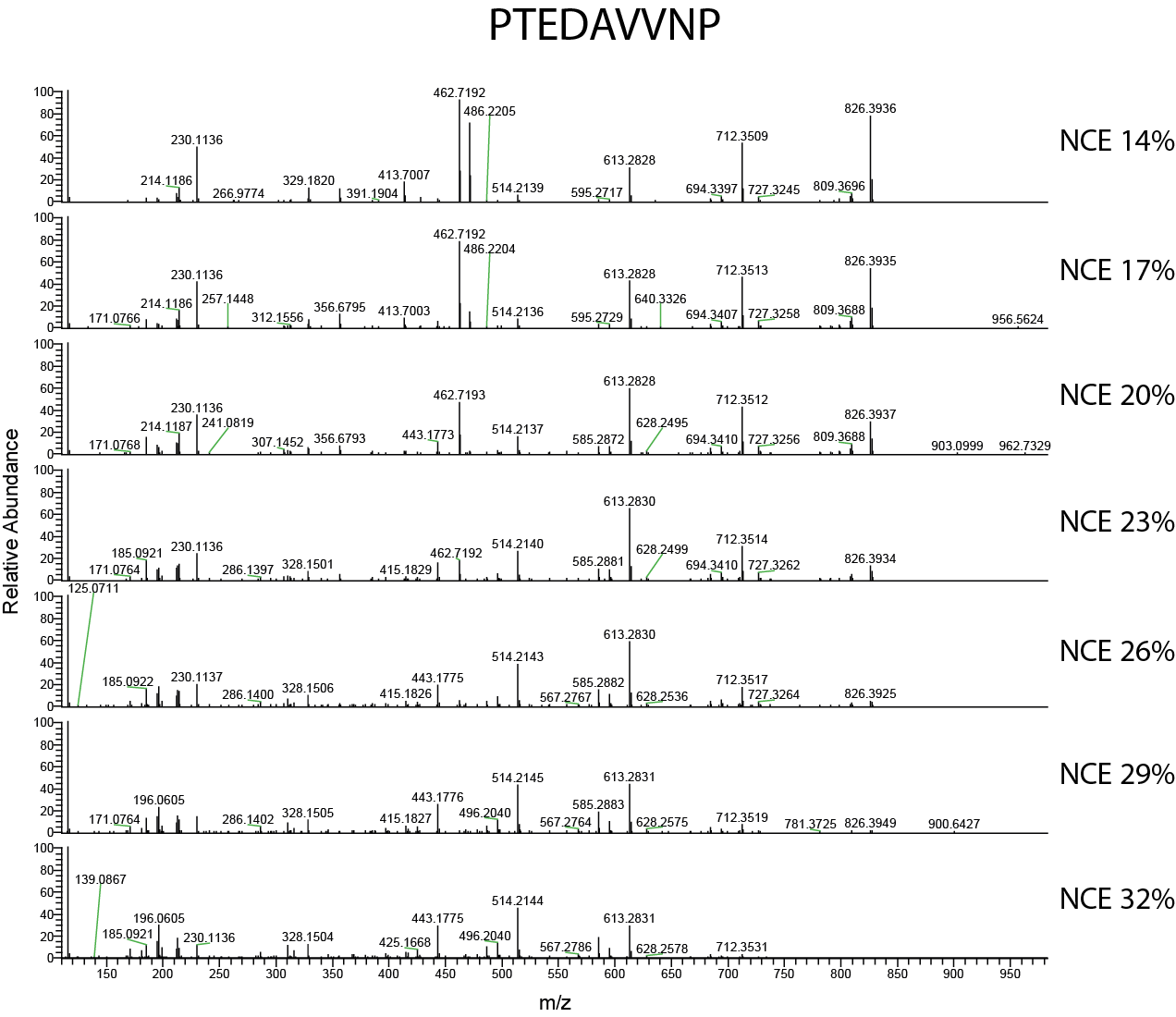

Figure S2. MS/MS fragmentation patterns at different collision energies**.** The peptide PTEDAVVNP was fragmented at several normalized collision energies (NCEs) during LC-MS/MS analysis. The b_7_ (m/z 712.351) and y_2_ (m/z 230.114) ions are essential to correctly assign the peptides.

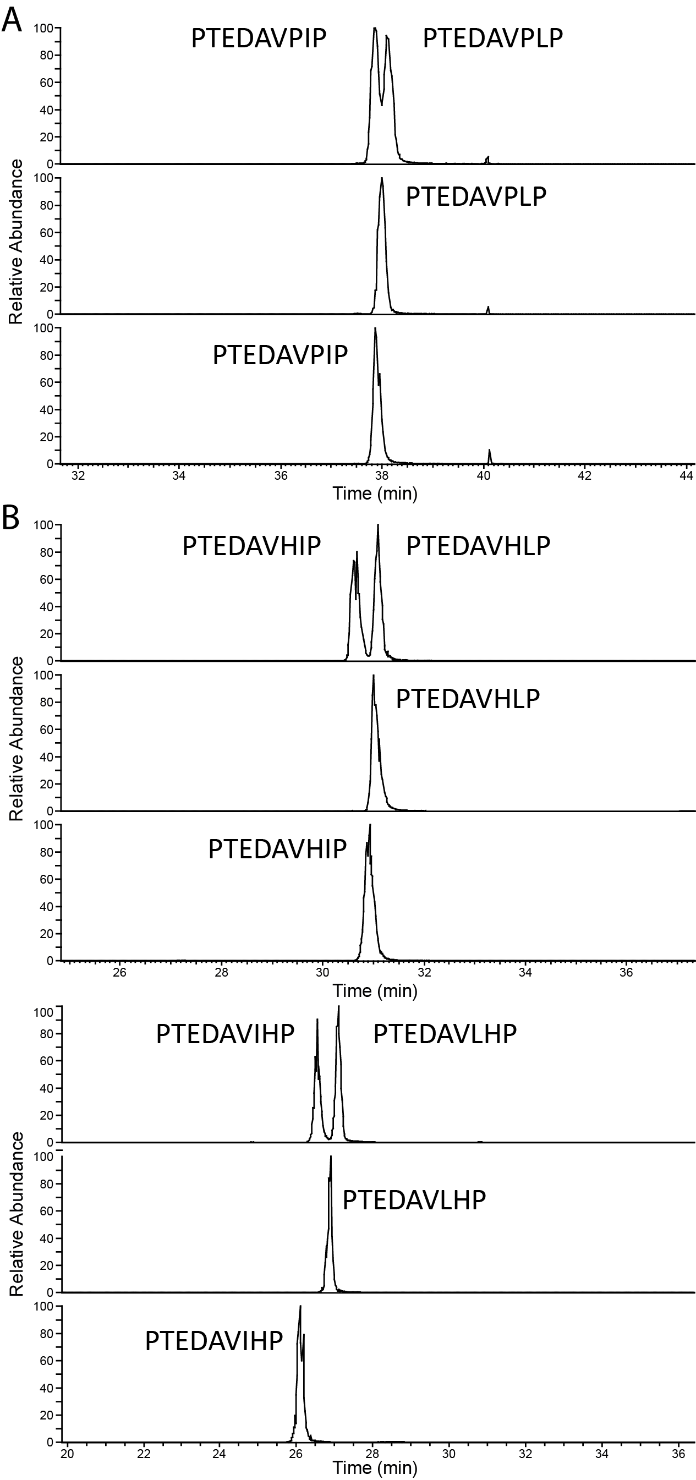

Figure S3. Chromatographic separation on a C18 column of synthetic peptides**.** To correctly assign product peptides with equal masses, several synthetic peptides were synthesized to assess the separation during LC. **A)** Separation of PTEDAVPLP and PTEDAVPIP. **B)** Separation of peptides from the non-prime-side library containing HIP/HLP/IHP/HLP (P3-P1).

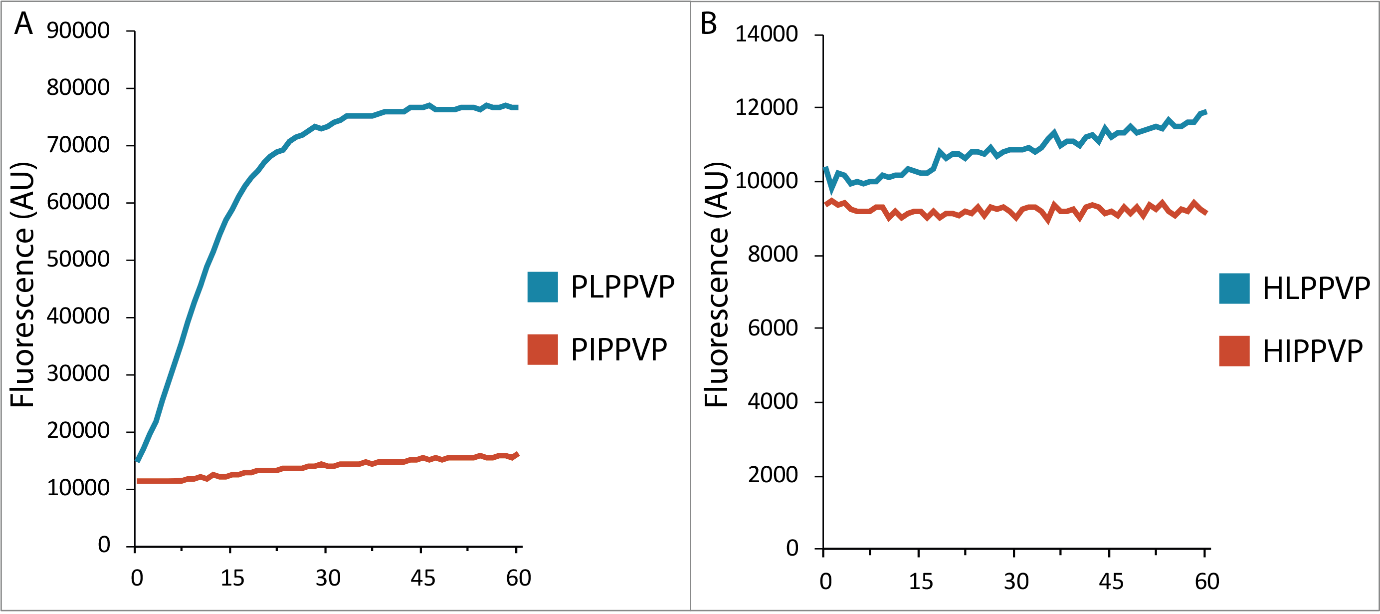

Figure S4. Cleavage of FRET-quenched peptides by PPEP-2**. A)** Time course of PPEP-2 mediated cleavage of the synthetic FRET-quenched peptides Lys(Dabcyl)-EP(I/L)PPVPD-Glu(EDANS). **B)** Time course of PPEP-2 mediated cleavage of the synthetic FRET-quenched peptides Lys(Dabcyl)-EH(I/L)PPVPD-Glu(EDANS).

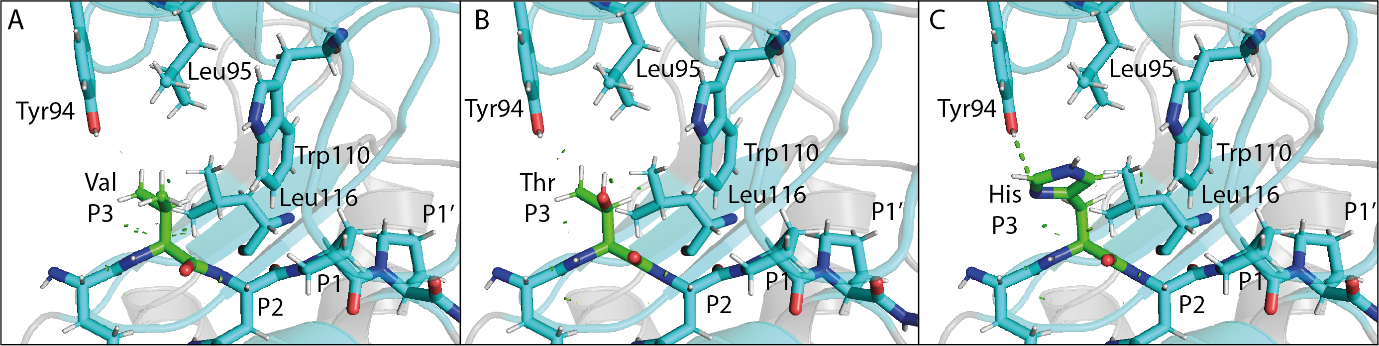

Figure S5. Modeled substitution of the Val at the P3 position in the PPEP-1 cocrystal with substrate VNPPVP**. A)** The cocrystal of PPEP-1 with substrate VNPPVP (*cyan*, PDB: 6R5C). The Val at the P3 position of the substrate is shown in *green*. The P3 contacting residues Tyr94, Leu95, Trp110, and Leu116 are shown as sticks. **B)** Substitution of the Val at the P3 position with Thr. No steric clashes or polar interactions were formed. **C)** Substitution of the Val at the P3 position with His. Polar interactions are shown as a green dotted line. Of note, a second rotamer in which the imidazole ring is rotated 180° is possible, but this produces a weaker interaction (hydrogen bonding distance=3.5 Å).

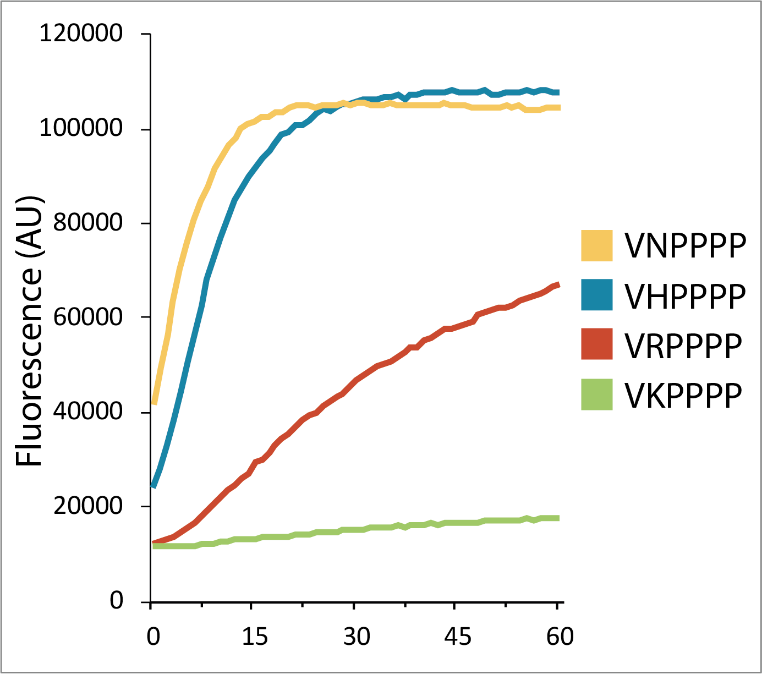

Figure S6. Cleavage of peptides containing Asn or the basic residues His, Arg, and Lys at the P2 position by PPEP-1**.** Time course of PPEP-1 mediated cleavage of the synthetic FRET-quenched peptides Lys(Dabcyl)-EV(V/H/R/K)PPPPD-Glu(EDANS).

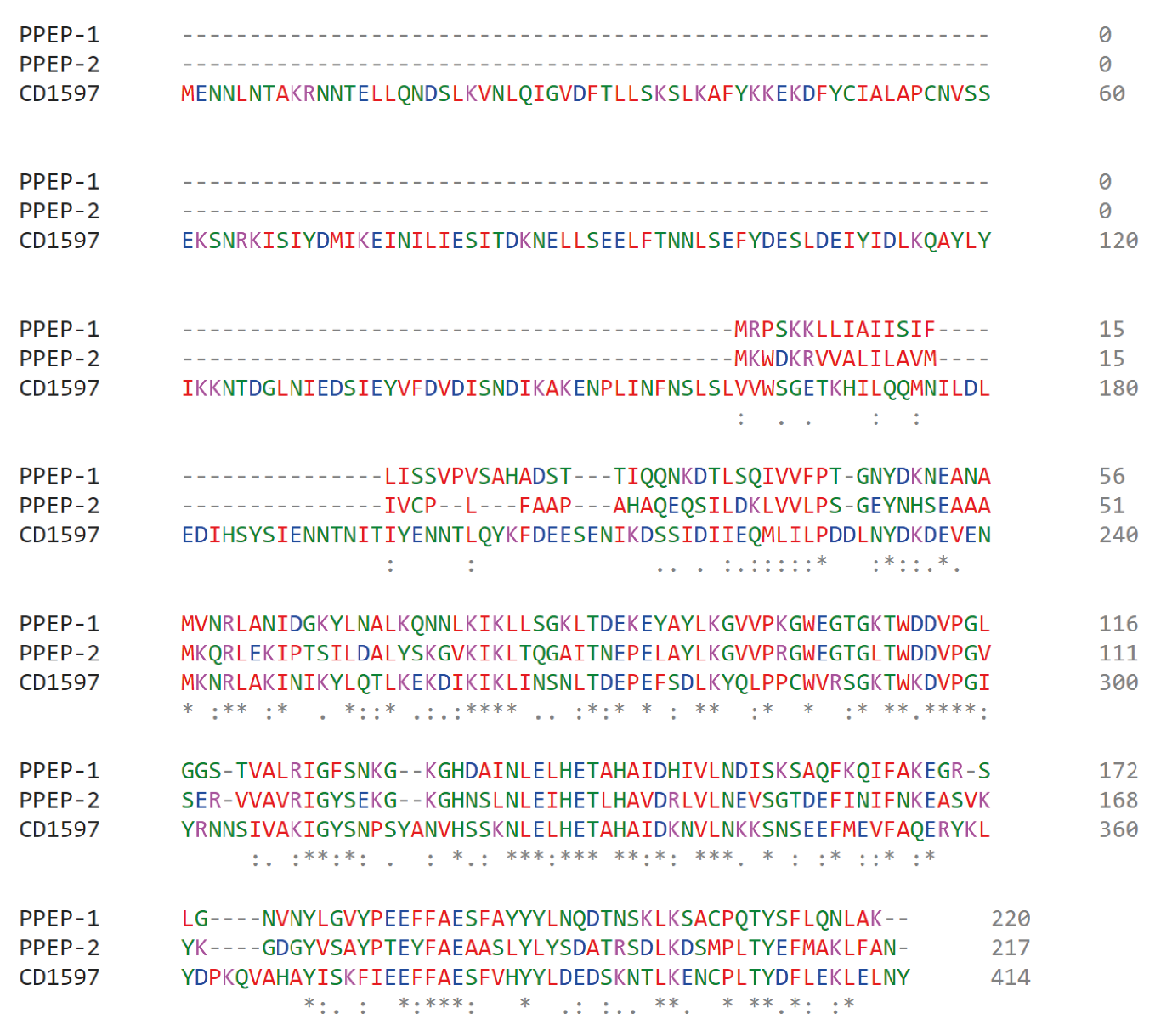

Figure S7. Sequence alignment of PPEP-1, PPEP-2, and CD1597**.** The sequence alignment was generated using the Clustal Omega Multiple Sequence Alignment tool.

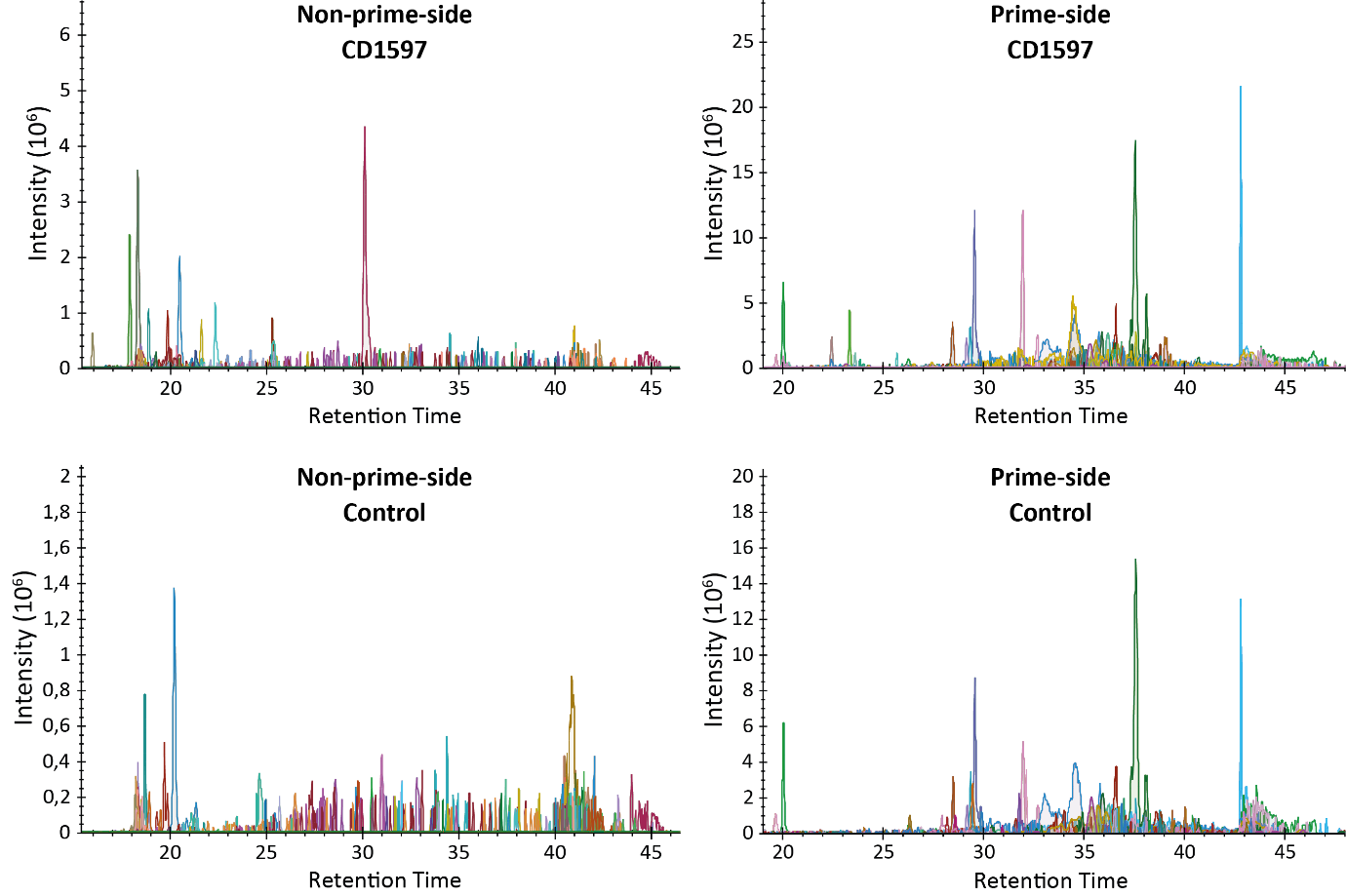

Figure S8. Incubation of the separate non-prime- and prime-side libraries with CD1597**.** The non-prime- and prime-side libraries were separately incubated with CD1597. EICs were constructed that include all possible 9-mer product peptides (PTEDAVXXP and PXXGLEEF).

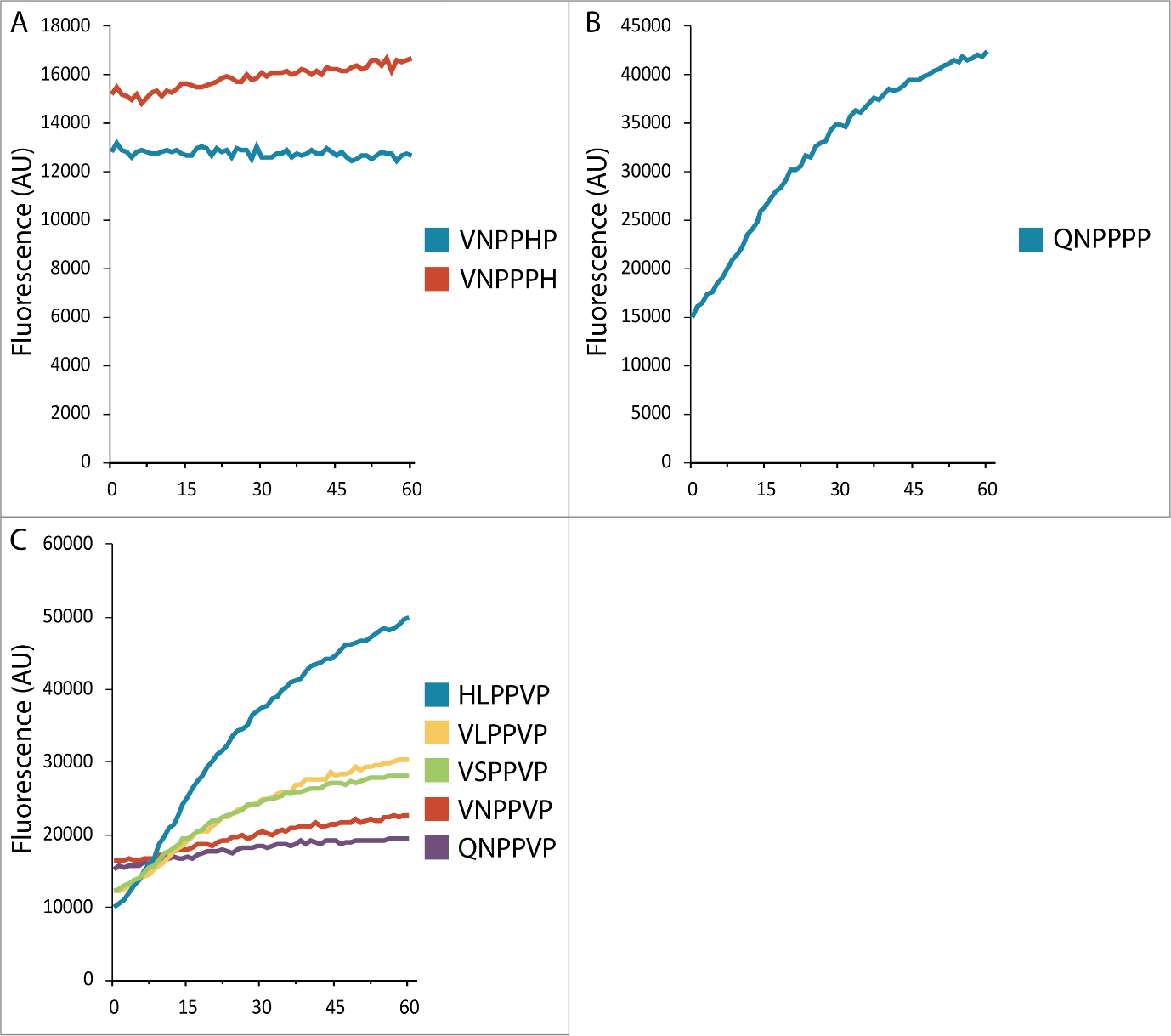

Figure S9. Cleavage of FRET-quenched peptides by PPEP-4**. A)** Time course of PPEP-4 mediated cleavage of the synthetic FRET-quenched peptides Lys(Dabcyl)-EVNPPHPD-Glu(EDANS) and Lys(Dabcyl)-EVNPPPHD-Glu(EDANS). **B)** Time course of PPEP-4 mediated cleavage of the synthetic FRET-quenched peptide Lys(Dabcyl)-EQNPPPP-Glu(EDANS). **C)** Time course of PPEP-4 mediated cleavage of the synthetic FRET-quenched peptides Lys(Dabcyl)-E(QN/VN/VL/HL/VS)PPVP-Glu(EDANS).

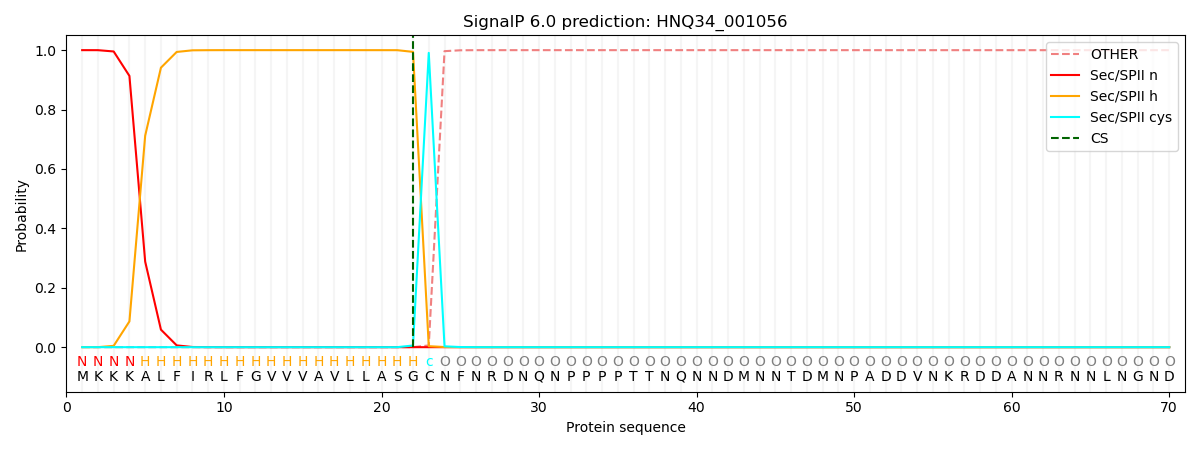

Figure S10. The predicted lipoprotein with a putative PPEP-4 cleavage site (gene: HNQ34_001056, UniProt ID: A0A7W8IQP4) from A. tepidamans is predicted to be inserted in the lipid membrane**.** Signal peptide cleavage occurs N-terminal of the cysteine as indicated by the blue “c”. Signal peptide prediction was performed using SignalP 6.0.
